# A Discard-and-Restart MD algorithm for the sampling of realistic protein transition states and enhance structure-based drug discovery

**DOI:** 10.1101/2024.06.14.598892

**Authors:** Alan Ianeselli, Jonathon Howard, Mark B. Gerstein

## Abstract

We introduce a Discard-and-Restart molecular dynamics (MD) algorithm tailored for the sampling of realistic protein transition states. It aids computational structure-based drug discovery by reducing the simulation times to compute transition pathways by up to 2000x. The algorithm iteratively performs short MD simulations and measures their proximity to a target state via a collective variable (CV) loss, which can be defined in a flexible fashion, locally or globally. Using the loss, if the trajectory proceeds toward the target, the MD simulation continues. Otherwise, it is discarded and a new MD simulation is restarted, with new initial velocities randomly drawn from a Boltzmann distribution. The discard-and-restart algorithm demonstrates efficacy and atomistic accuracy in capturing the folding pathways in several contexts: (1) fast-folding small protein domains; (2) the folding intermediate of the prion protein PrP; and (3) the spontaneous partial unfolding of α-Tubulin, a crucial event for microtubule severing. During each iteration of the algorithm, we are able to perform AI-based analysis of the transitory conformations to find binding pockets, which could potentially represent druggable sites. Overall, our algorithm enables systematic and computationally efficient exploration of conformational landscapes, enhancing the design of ligands targeting dynamic protein states.

## INTRODUCTION

Structure-based drug discovery (SBDD) provides a rational and quantitative approach for the design of therapeutic ligands targeting specific protein structures involved in disease. It relies on the knowledge of the atomistic 3D structure of a biological macromolecule, which can be obtained by means of experimental (e.g. NMR, X-Ray) and computational (e.g. Molecular Dynamics, AlphaFold) techniques^1–4^. The 3D structure of the macromolecule is then used to design small molecules that bind specific regions with high affinity and selectivity.

For what concerns the computational aspects of SBDD, one of the greatest challenges is the computational time required for an accurate modeling of the conformations of biological macromolecules. MD simulations often require prohibitive computational times to simulate the dynamic behavior of proteins and their conformational transitions^5,6^, to identify druggable regions of the macromolecule and design structure-specific drug ligands. There is, therefore, a pressing need for innovative computational approaches to accelerate the SBDD computational workflow, yet maintaining the atomistic accuracy required for a successful predictive model.

In this paper, we propose a new algorithm to generate trajectories of conformational transitions via plain MD. It relies on the description of a target state via a collective variable (CV) loss function. Short MD simulations are iteratively performed and restarted-and-discarded or continued based on their proximity to the target state, described by the CV. We have tested it on well-characterized fast-folding proteins. The algorithm has accurately computed the key features of their folding and many atomistic characteristics, but at a fraction of the computational cost that would be required by unguided MD. We have also applied it to study the folding intermediate of the prion protein and the spontaneous unfolding process of α-Tubulin. Even without an explicit definition of the target state (the spontaneous partially unfolded state of α-Tubulin is unknown), the method was able to generate its unfolding trajectory and obtain the transition states at atomistic detail. With the use of AI tools, we have predicted the pocket binding sites of each transition state and designed drug ligands able to dock them with high affinity.

Our algorithm yields a computationally efficient way to compute the transition states of biomolecules, allowing for the identification of realistic dynamic protein states to be used in the workflow for structure-based drug discovery studies.

### Discard-and-restart algorithm

The algorithm is shown in Figure 1. Starting from an arbitrary protein conformation, a short (10-20 ps) MD simulation is run. The trajectory is measured via collective variables which describe the conformational changes. The choice of CVs can be optimized and varied depending on the problem to be investigated. If the new conformation’s CV loss is lower, the conformation is accepted and will be used as the new starting conformation for the next simulation. On the other case, if the loss increases, the simulation is discarded and restarted using a different random seed for the initial atomic velocities.

**Figure 1.**
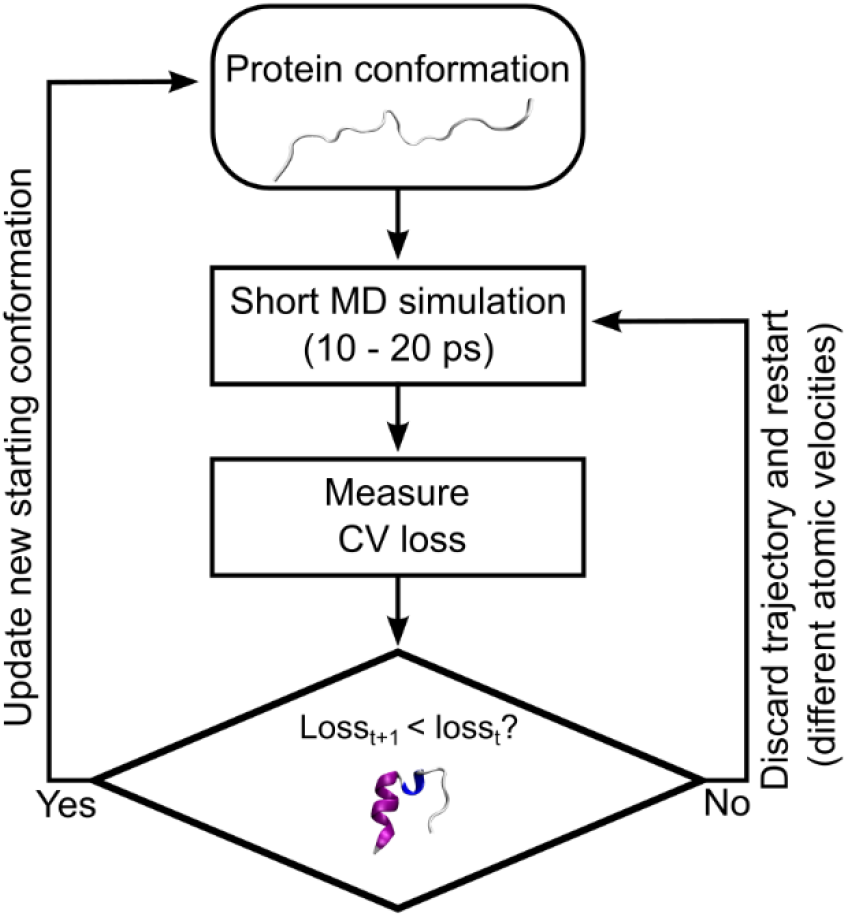
The discard-and-restart algorithm.

In the MD simulations (using GROMACS^7^), the initial velocities of the atoms were generated from a Maxwell-Boltzmann distribution at a given temperature (the thermostat temperature). Each new short MD simulation, therefore, starts with different initial velocities, leading to a diverse exploration of the conformational space. This is a similar approach to Monte Carlo (MC) simulations^8^, where new conformations are obtained by performing dihedral torsions, bond angle movements and bond stretchings. However, it is important to note that in MC simulations, the system explores the conformational space via stochastic moves. In contrast, in our algorithm, MD simulations evolve the system deterministically according to Newton’s equations of motion, although the starting conditions for each trajectory (i.e., the initial atomic velocities) may be drawn from the Maxwell-Boltzmann distribution.

Additionally, if the trajectory is unable to go forward for too many times, the new starting conformation is updated regardless, using the last frame of the simulation. This is to allow the protein to exit local energy minima it could get stuck into. More details are in supplementary section 1-2. Note that we have used the MD engine (GROMACS) as it is, without any under-the-hood modifications and without the addition of any biasing forces to the protein atoms or modifications of the energy landscape of the protein. We have modeled the proteins using the Amber99SB-ILDN force field^9^.

The idea behind this algorithm comes from a primordial DNA sequence evolution paradigm in the context of the origin of life, where DNA sequences that are able to replicate survive and continue, while those who are not fit stall and die out^10,11^.

Our algorithm shares similarities with other ones such as milestoning^12^, weighted ensemble^13^, forward-flux sampling^14^, RESTART approach^15^. However, these methods require prior knowledge of the system such as the energy landscape, the structure of the intermediate conformations, or the position of the thresholds of the transition states. In our case, no such prior knowledge is required. The only pre-requisite is the choice of a collective variable (CV) that can best describe the system and its progression towards the target. Moreover, as we will see in the section regarding α-Tubulin, the target does not need to be explicitly defined.

## RESULTS

### Folding of small protein domains

#### Collective variable for protein folding

The process of protein folding is often described through one or multiple collective variables. We have identified four CVs which, according to past and recent literature, are able to describe the fine details of the folding process and have, in fact, also been employed during metadynamics^16^, ratchet-and-pawl MD^17^, Greedy-Proximal^18^ and Weighted Ensemble^19^ algorithms. These four CVs are described in the following lines and are shown in Figure 2a-d:

**Figure 2.**
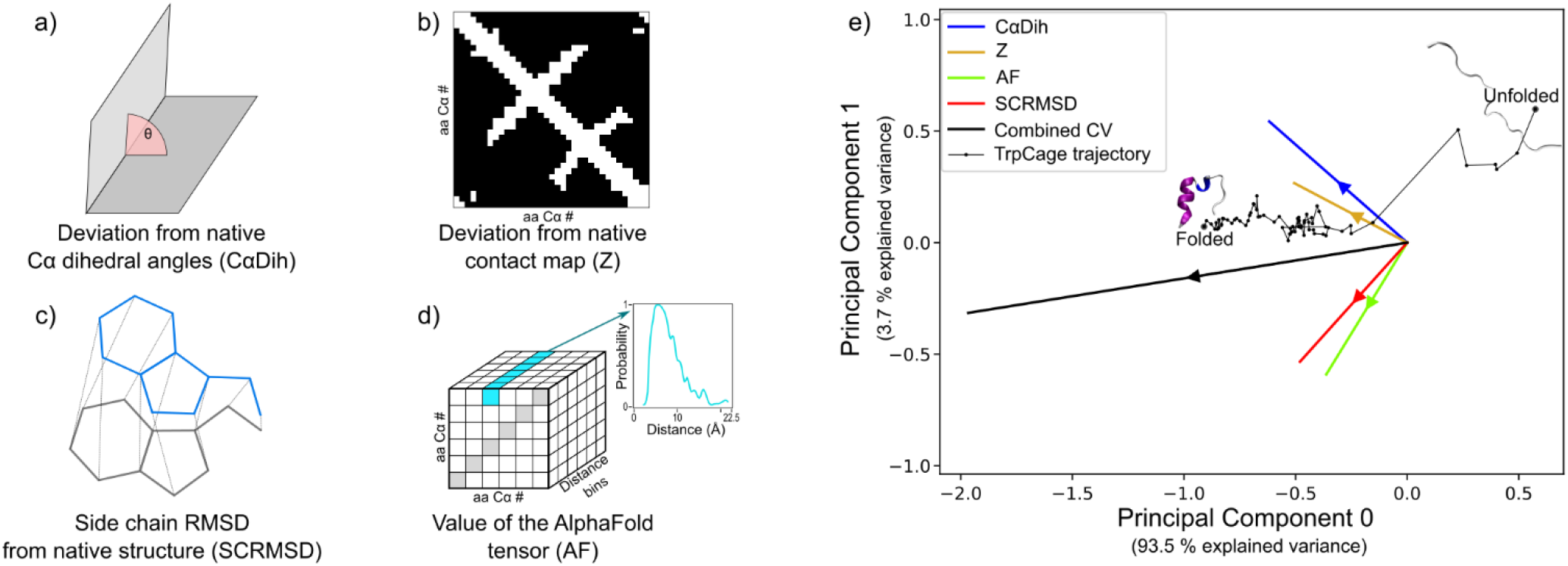
Global CV for protein folding. Four CVs are particularly able to describe the fine details of protein folding: (a) deviation of Cα dihedrals from the native state ones; (b) deviation from the native Cα contact map; (c) RMSD of the side chains from the native structure; (d) the value of the tensor derived from Google’s Deepmind Alphafold 2 AI tool. e) The four CVs have been combined by PCA, calculating the PCA eigenvectors on long MD trajectories of the Villin subdomain and Fip35 WW domain provided by DE Shaw Research. The combined CV is then used with the discard-and-restart algorithm to determine whether the trajectory is moving toward the target (native) state or not. The black thin dotted line shows a TrpCage miniprotein folding trajectory onto the PCA plane.

- Deviation of Cα dihedrals from the native state (CαDih) 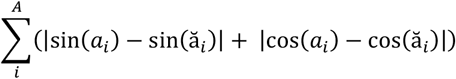

where *A* is the total number of Cα atoms in the protein, *a*_*i*_ and *ă*_*i*_ correspond to the *i*^th^ dihedral angle of one conformation and the native state, respectively, calculated on Cα atoms only. This CV quantifies the difference in the torsion of the protein backbone between two conformations.
- Deviation from the native Cα contact map (Z) 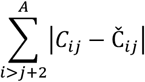

where *A* is the total number of Cα atoms in the protein, *C*_*ij*_ and *Č*_*ij*_ are the contact map entries of one conformation and the native state, respectively, and are defined as: 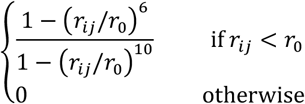

where *r*_*ij*_ is the Euclidean distance between atom *i* and *j, r*_*0*_ is a conventional reference contact distance for contact formation set to 0.75 nm. The constraint |*i*>*j*+2| in the summation ensures that consecutive contacts are excluded from the calculation. This CV quantifies the deviation of the protein contacts (Cα atoms closer than 0.75 nm) from the target (i.e. native) conformation. The value of Z decreases whether when native contacts are formed or non-native contacts are disrupted.
- Side chain RMSD from the native state (SCRMSD) 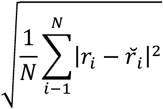

where *r*_*i*_ and *ř*_*i*_ are the Cartesian coordinates of the *i*^th^ side chain atom (excluding hydrogens) of one conformation and the native state (after alignment), respectively, and *N* is the total number of atoms. This CV quantifies the geometrical deviation between the side chains of two protein conformations.
- Value of the AlphaFold tensor (AF) One of the outputs of Google Deepmind AlphaFold 2, an outstanding AI milestone for the prediction of protein structures from their aa sequence^3^, is a tensor *D* of distance logits with shape *A×A*×*M*, where *A* is the number of aa in the protein and *M* is the number of distance bins (64 bins ranging from 0.23 to 2.17 nm). The tensors have been calculated using AlphaFold 2 on the COSMIC2 server platform^20^. After converting logits to probabilities (exp(*D*)/(1+exp(*D*))), the AlphaFold-derived CV (AF) was defined as following^16^:

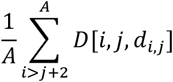

where *d*_*i*,*j*_ corresponds to the index of the distance bin where the distance of the residues *i*,*j* falls within. Again, the constraint |*i*>*j*+2| in the summation ensures that consecutive contacts are excluded from the calculation. This CV is a probabilistic evaluation of how a conformation would be favored by the AlphaFold 2 model. As shown in Figure 2d, *D*[*i*,*j*] is a vector generated by AlphaFold 2 that indicates the probability to find the residues *i* and *j* at a given distance. Being generated by an AI model trained on ∼170k protein structures, it can be interpreted as the quasi-chemical energy potential between amino acid pairs and relates to the strength of such pairwise interactions^21,22^.

### CV loss for protein folding

To unequivocally assess whether the protein is moving towards the target state or not, it is necessary to have a single, one-dimensional CV that can best describe the protein dynamics. We have adopted the strategy to merge the four CVs discussed above into one via Principal Component Analysis (PCA)^23^. The PCA eigenvectors have been calculated using very long plain MD simulations of Villin and Fip35 WW domain (100 µs for each protein), kindly provided by DE Shaw Research. More details in the supplementary information, section 1. Results are show in Figure 2e. Hence, it was possible to calculate the one-dimensional, combined CV during our simulations for any protein conformation (e.g. Figure 2e, black thin line). This combined CV of reduced dimensionality could be considered as an optimized CV for the folding of small proteins. It is a generalized measure that describes the global conformational changes of small protein domains.

### Folding trajectories of small protein domains

We have benchmarked our algorithm on four fast-folding small protein domains whose folding pathway has been characterized several times by plain MD simulations and other methods^18,24–26^: TrpCage miniprotein (PDB: 1L2Y, 20 aa); Fip35 WW domain (PDB: 2F21, 35 aa); Villin subdomain (PDB: 2F4K, 35 aa); β-hairpin fragment (PDB: 1gb1, 16 aa). Results are shown in Figure 3.

**Figure 3.**
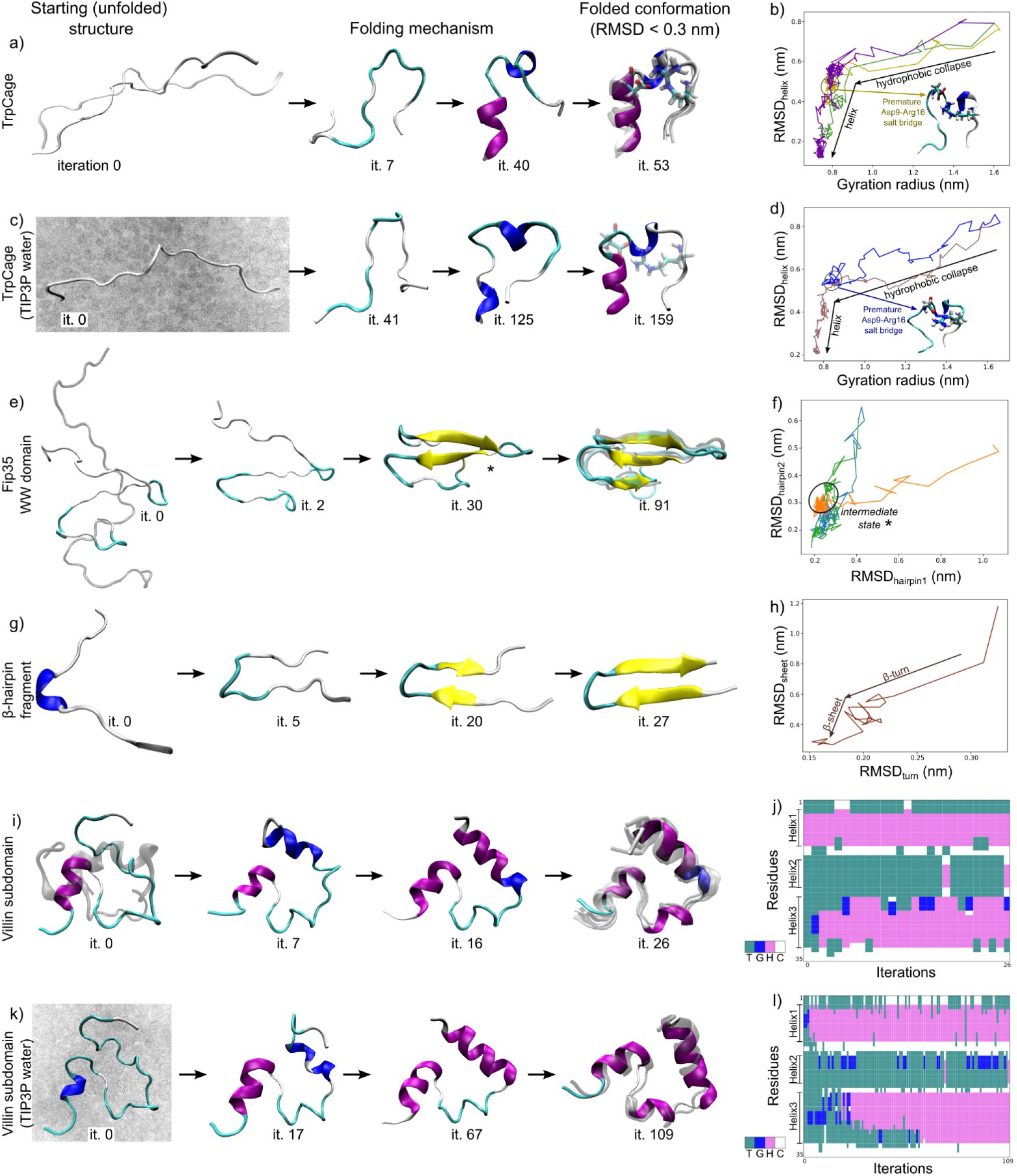
Folding of small protein domains by Discard-and-Restart MD. (a) TrpCage folding and RMSD_helix_ vs gyration radius plot (b). The protein undergoes an initial hydrophobic collapse, followed by the formation of the α-helix and backbone rearrangements, stabilized by a Asp9-Arg16 salt bridge. The trajectory gets stuck in an intermediate state when the salt bridge is formed prematurely. (c) TrpCage folding in explicit solvent (TIP3P water), and RMSD_helix_ vs gyration radius plot (d). The folding mechanism is qualitatively identical to the one obtained in implicit solvent. (e) Fip35 WW domain folding, and RMSD_hairpin1_ vs RMSD_hairpin2_ plot. (f). Hairpin 1 folds before hairpin 2. An intermediate with a native-like hairpin 1 but an unstructured hairpin 2 has been observed. (g) β-hairpin folding and RMSD_turn_ vs RMSD_sheet_ plot (h). The core (β-turn) forms first and is then followed by the progressive formation of the β-sheet. (i) Villin folding in implicit solvent and its DSSP plot during the iterations (j). Helix 1 and helix 3 form first, followed by helix 2 which requires many more iterations. (k) Villin folding in explicit solvent (TIP3P water) and DSSP plot (l). The folding mechanism is qualitatively the same as the one obtained in implicit solvent. DSSP plot legend: turn (T); 3-10 helix (G); α-helix (H); coil (C).

#### TrpCage miniprotein (implicit solvent)

The starting unfolded structures have been taken from the repository of the paper by Spiwok *et al*^16^, which they have used as starting structures in their metadynamics simulations. We performed a total of 12 simulations at 300 K starting from 3 different initial configurations. 2 trajectories reached the native state after 58 and 143 iterations (fastest folding time 254 ps).

According to past computational and experimental studies, the folding mechanism of the TrpCage miniprotein mostly consists in two pathways^27–32^. Pathway I, in which the hydrophobic collapse of the core precedes the formation of the α-helix. Pathway II, in which the events occur in the opposite order, i.e. the α-helix forms first followed by the hydrophobic collapse. Pathway I prevails in simulations at room T, while pathway II becomes dominant at melting and higher temperatures.

In our simulations, all TrpCage trajectories followed the pathway I: from the unfolded conformation, the core of the protein underwent a rapid collapse, and then the formation of the α-helix followed (Figure 3a-b). We have not observed pathway II in our simulations, and this is probably due to the fact that the simulations have been run at 300 K, a temperature where pathway I is dominant. We have also observed the formation of the salt bridge between Asp9 and Arg16, which is thought to play an important role in the stabilization of the folded state^33,34^. We have also observed a conformation resembling the intermediate state reported in the literature^35,36^, in which some of our trajectories got stuck (Figure 3b inset). This state was characterized by the presence of a premature salt bridge between Asp9 and Arg16, prior to the formation of the α-helix. It is thought that the premature formation of the Asp9-Arg16 salt bridge blocks the protein in an intermediate metastable state, because the salt bridge needs to be broken for the next native rearrangements to take place. This therefore creates a high activation energy barrier that prevents this conformation from folding^35^.

#### TrpCage miniprotein (explicit solvent)

The starting unfolded structure wastaken from the repository of the paper by Spiwok *et al*^16^. Out of a total of 5 simulations (temperature 300 K), 1 reached the native state (RMSD < 0.3 nm) in 159 iterations (595 ps).

The folding pathway of this trajectory was qualitatively identical as the one obtained in implicit solvent, with an initial hydrophobic collapse of the structure followed by the formation of the helix and the rearrangements of the backbone (Figure 3c-d). The salt bridge between Asp9-Arg16 also formed in the native state and further stabilized the native. Again, we have observed in another trajectory that the premature formation of the salt bridge gets the protein stuck in an intermediate metastable state, blocking further native rearrangements from taking place (Figure 3d inset).

#### Fip35 WW domain

The starting unfolded conformations have been generated by thermal unfolding at 700 K. Out of a total of 16 simulations (run at 300 K or 350 K in implicit solvent) starting from 2 different unfolded configurations, 3 folded (RMSD < 0.3 nm) after 67 to 91 iterations (fastest folding time 332 ps).

Past studies indicate that the folding pathway of Fip35 WW domain proceeds in a very heterogeneous way, through a multitude of qualitatively different but equiprobable folding pathways, which mostly differ by the order of formation of the two hairpins^37,38^. In our case, all folding trajectories followed the pathway in which hairpin 1 folds before hairpin 2 (Figure 3e-f). This is probably due to the low number of trajectories and initial conditions that we have simulated, which limited the explored conformational space. Interestingly, our simulations showed the existence of a long-lived intermediate state with hairpin 1 in a native-like configuration and hairpin 2 still unstructured (Figure 3e, third snapshot). This observation is in agreement with past MD trajectories of the Fip35 WW domain^39^, and can also be seen in the long MD trajectories of DE Shaw Research^40^.

#### β-hairpin fragment

The starting unfolded structure has been generated via thermal unfolding at 700 K in implicit solvent. One trajectory (out of a total of 3 simulations at 325 K) reached the native conformation (RMSD < 0.3 nm) after 27 iterations (folding time 166 ps). The folding mechanism obtained in our simulations (Figure 3e-f) is in agreement with what is reported in the literature: the folding of the hairpin occurs by first forming the core (the β-turn) and then β-sheet hydrogen bonds, which progressively propagate along the strands^25,41,42^.

#### Villin subdomain (implicit solvent)

The starting unfolded structures have been taken from SimTK in the repository associated with the paper by Ensign et al^43^. These structures had been generated by long MD simulations of thermal unfolding at 373 K. We have chosen the conformations that folded the fastest in their study (structures 4 and 7). In our simulations at 300 K, out of a total of 22 simulations, 7 reached the native state (RMSD < 0.3 nm) after 20 to 29 iterations (fastest folding time 134 ps).

According to past studies, the folding mechanism of villin is very complex and heterogeneous^43–45^, and the pathways strongly depend on the temperature, on the starting configuration and on the type of force field model chosen^43,46^. In our simulations, we have observed that the first secondary structures that form are helix1 and helix3, which form simultaneously (Figure 3i-j). Helix2 starts to form later and requires more iterations than helix1 and helix3 in order to fully form. This observation is in agreement with a recent study^47^, which classifies villin’s folding pathways based on the mechanism of formation of helix2, after helix1 and helix3 are already formed.

However, due to the low number of simulations and initial configurations, we could not successfully sample the heterogeneity of the villin’s folding space. Moreover, the folding mechanisms that we observed were probably influenced by the fact that the initial configurations from Ensign *et al*^43^ were pretty compact and helix1 and helix3 were not completely unfolded.

#### Villin subdomain (explicit solvent)

The starting unfolded structure was taken from SimTK^43^ (structure 4), as indicated in the previous section. Out of a total of 6 simulation (run at 300 K), 2 reached the native state (RMSD < 0.3 nm) after 109 and 119 iterations (fastest folding time 586 ps).

The folding pathway of the folding trajectories was qualitatively the same as the one obtained in implicit solvent (Figure 3k-l). Helix1 and helix 3 start to form first, almost simultaneously, followed by helix 2 which requires many more iterations. Also in this case, the number of initial conditions and simulations that we performed is very limited and did not allow us to explore the complex and heterogeneous folding landscape of the villin subdomain.

Overall, only a fraction of the simulations that we have performed reached the folded state (∼33%). Many trajectories got stuck in local energy minima, and could not go forward anymore, even though they formed local secondary structures and reached a partial native-like conformation. It is possible that they could have managed to jump out and continue their path towards the folded state, given a sufficient number of attempts and simulation duration. Moreover, a better tuning of the algorithm and MD parameters could have increased the number of successful folding trajectories. For example, increasing (or decreasing) the simulation length of each iteration, modifying the value of the *Rappa*, changing the temperature of the MD simulation, or choosing better CVs. Table 1 summarizes the results obtained for the fast-folding proteins.

**Table 1.** Summary of folding results for the fast-folding proteins.

| Protein | Size (aa) | PDB | # starting structures | Solvent model | T (K) | Total n° of simulations | n° of folding | Folding efficiency | Fastest folding time | Avg CPU time per folding traj |
| --- | --- | --- | --- | --- | --- | --- | --- | --- | --- | --- |
| TrpCage miniprotein | 20 | 1L2Y | 2 | Implicit (GBSA) | 300 | 12 | 3 | 25 % | 254 ps | ~18 h * 1 core =18 h |
| TrpCage miniprotein | 20 | 1L2Y | 1 | Explicit (TIP3P) | 300 | 5 | 1 | 20 % | 1190 ps | ~164 h * 6 core =984 h |
| Fip35 WW domain | 35 | 2F21 | 2 | Implicit (GBSA) | 300<br>350 | 16 | 3 | 19 % | 332 ps | ~39 h * 2 cores =78 h |
| β-hairpin fragment | 16 | 1gb1 | 1 | Implicit (GBSA) | 325 | 3 | 1 | 33 % | 166 ps | ~1.5 h * 1 core =1.5 h |
| Villin subdomain | 35 | 2F4K | 2 | Implicit (GBSA) | 300 | 22 | 7 | 32 % | 134 ps | ~38 h * 4 cores =152 h |
| Villin subdomain | 35 | 2F4K | 1 | Explicit (TIP3P) | 300 | 6 | 2 | 33 % | 586 ps | ~90 h * 4 cores =360 h |

We have compared the CPU times to fold the fast-folding proteins by unguided MD vs our Discard-and-Restart MD algorithm. Results are shown in supplementary section 3. It reduces the computational CPU times to achieve folding of a factor up to ∼2000x (range between 20x to 2000x, average 510x).

## RESULTS

### Folding intermediate of PrP

The cellular prion protein (PrP) is a cell surface glycoprotein that is involved in numerous regulatory cellular pathways. PrP is also responsible for the transmissible spongiform encephalopathies, caused by its conformational conversion into an aggregated pathogenic form (PrP^Sc^) which creates protein aggregates in the brain and disrupt normal brain functions^48,49^. The study of its folding pathway and conformational interconversion is therefore of great therapeutic interest.

Numerous attempts to pharmacological target the PrP native state have failed, suggesting that the native conformation could be an undruggable target^50,51^. With the aim of targeting a folding intermediate pharmacologically, a recent study has fully characterized its folding pathway and identified a folding intermediate which was successfully targeted by a specific drug ligand^52^. PrP presents a native-like folding intermediate where helix 1 (H1) is displaced and misses its docking contacts with helix 3 (H3), exposing to the solvent several residues that, in the native state, would be buried inside the protein core. This was the first study to characterize PrP’s folding pathway at an atomistic level. Other experimental studies indicated the presence of a native-like intermediate state with congruent structural characteristics^53–56^.

### CV loss for PrP folding intermediate

The PrP intermediate is characterized by H1 displaced from H3. The CV to study the transition therefore is simply a local measure of the distance between the helix H1 and the helices at the opposite side of the protein end (H2H3) (Figure 4a):

**Figure 4.**
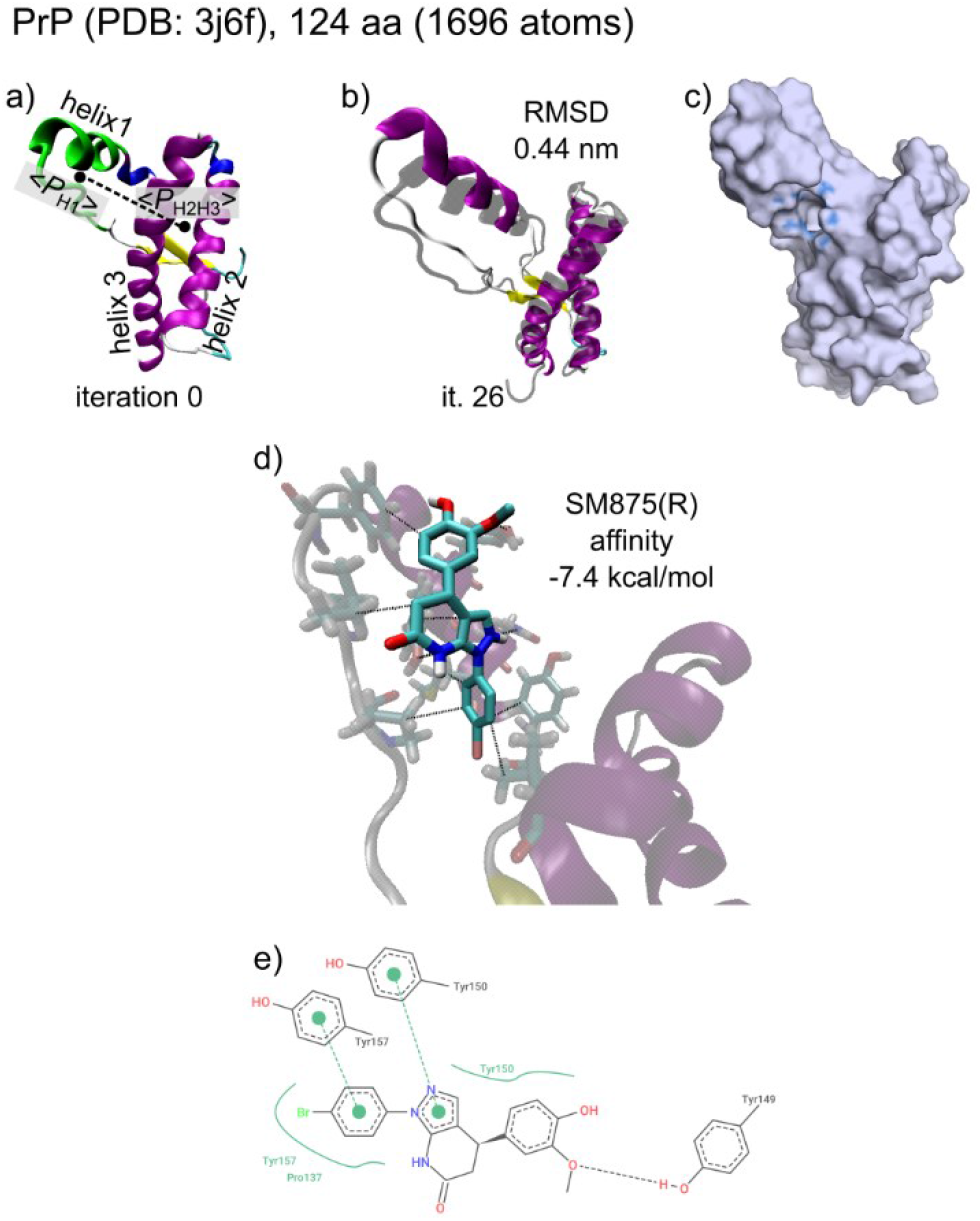
Transition of PrP from native to intermediate. a) Structure of the native PrP (C-terminal structured region) and illustration of the CV. The large dots indicate the average coordinate point for H1 and H2-H3. b) Structure of the intermediate conformation obtained by Discard-and-Restarted MD superimposed to the original intermediate state from Spagnolli *et al*. (gray transparent structure). c) Surface image and ligand binding site predicted with P2Rank (blue). d-e) Predicted ligand binding pose for SM875(R) docked onto the intermediate druggable pocket shown in (c).

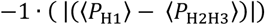

where <*P*> corresponds to the average position of the atoms of helix 1 (H1, residues 138-153) and helices 2 and 3 (H2-H3, residues 174-226).

### Simulation of PrP intermediate transition

The starting structure has been the native, structured C-terminal domain of the human PrP protein (residues 125-228, 1696 atoms, PDB: 1QLX). We have simulated the transition between the native structure and the intermediate, as the transition is reversible and therefore also accessible from the native state. The simulations have been run at 350 K, for a total of two trajectories. Each simulation required ∼50 CPU hours.

In 26 iterations, the H1 progressively displaced from H2-H3 and lost the docking interactions with the protein core. The conformational change is slight, but large enough to expose several residues that would otherwise be buried inside of the protein core. The conformation that we have obtained is substantially identical to the one identified by the previously mentioned study^52^ (RMSD 0.44 nm, Figure 4b).

### Drug discovery on the transition states

During the PrP conformational change to the intermediate state, a lot of protein regions that were previously buried became exposed to the solvent. We have explored the possibility that these regions could have become targetable by ligands. We have utilized the tool P2Rank^57^ in order to predict the ligand binding sites of the transitory conformations. P2Rank is a machine-learning based tool that was trained on various structural and physicochemical features of proteins with known ligand binding sites. Given a protein structure, P2Rank predicts and ranks solvent-accessible protein regions according to their ligandability score. Figure 4c shows the ligand binding site predicted for the newly exposed surface region of the transition state, which involves the residues 137, 150, 153, 154, 157, 205, 209. Our pocket is just a few Angstrom away from the ligand binding site identified in the study from Spagnolli *et al*^52^ and shares most of the residues.

We have docked the ligand SM875(R), the drug molecule developed for the targeting of the PrP intermediate state, onto our pocket. We have used the docking tool Autodock Vina^58^ available on the SeamDock web server^59^ (https://bioserv.rpbs.univ-paris-diderot.fr/services/SeamDock/) or Webina^60^ (https://durrantlab.pitt.edu/webina/). It estimates the pose and affinity between a ligand and a receptor based on Van der Waals interactions, hydrogen bonding, electrostatic interactions and torsional terms. We have generated a squared cube of 2 × 2 × 2 nm around our binding pocket.

The docking simulation predicted a highest affinity pose of -7.4 kcal/mol, which is a strong interaction. The pose of the ligand in the pocket with the main protein-ligand interactions is shown in Figure 4d-e. Some interactions are the same as the ones in the study mentioned before (for example, the aromatic-aromatic interactions with Tyr157). A longer unfolding trajectory, with a similar intermediate state and ligand binding site is shown in the supplementary section 4.

## RESULTS

### Partial unfolding of α-tubulin helical domain H11-H11’-H12

Tubulin is a heterodimeric protein made of α- and β-subunits which assemble into microtubules, a fundamental component of the cytoskeleton of eukaryotic cells. The turnover of the microtubules is facilitated by severing enzymes, which include spastin and katanin^61–63^. These proteins are thought to pull tubulin subunits out of the microtubule lattice by pulling on their C-terminal tails (CTTs)^64^ and unfolding the peptide chain leading to removal of the tubulin subunit from the lattice. When enough tubulin subunits are removed, the microtubule filament breaks.

A recent study has shown experimentally that during the pulling of α-Tubulin out of the microtubule lattice, a partially unfolded, transitory intermediate is formed^65^. The authors of the study have hypothesized that the unfolding region involves the last helices of α-tubulin, namely H11-H11’-H12. Their kinetic calculations also indicate that this region can spontaneously unfold on the timescale of seconds, though in the absence of a pulling force the region refolds. However, in the presence of a pulling force, tubulin further unfolds leading to its removal from the lattice. Our goal is to simulate this transitory unfolding of the H11-H11’-H12 region. It is possible that when spastin pulls on the CTT it stabilizes the unfolded intermediate, facilitating the further unfolding of the tubulin peptide. The severing of microtubules by spastin may have mechanic similarities to the way actin filaments are severed by the enzyme cofilin^66,67^. Cofilin, in order to complete the binding to F-actin, needs to wait for spontaneous conformational fluctuations of the SD1 subdomain of F-actin. Only then it is able to fully bind to actin filaments and destabilize them to induce severing. It is therefore possible that conformational fluctuations in the H11-H11’-H12 domain play a similar role in microtubule severing by spastin.

α-tubulin (PDB: 3j6f) is a medium-sized protein (∼48 kDa) made of 428 aa (6608 atoms). Exploring its spontaneous partial unfolding process by unguided MD is computationally demanding because of the protein’s size and the timescale of its spontaneous unfolding. Our discard-and-restart MD algorithm has instead been able to capture this conformational event in reasonable computational times, as we will show in the next sections.

### CV loss for α-Tubulin unfolding

As discussed before, the partial unfolding event is likely to involve the helices present in the C-terminal region of α-Tubulin, in particular the helices H11, H11’ and H12. The CV for this transition has therefore been chosen to quantify the conformational changes in the C-terminal helical domain (Figure 5a):

**Figure 5.**
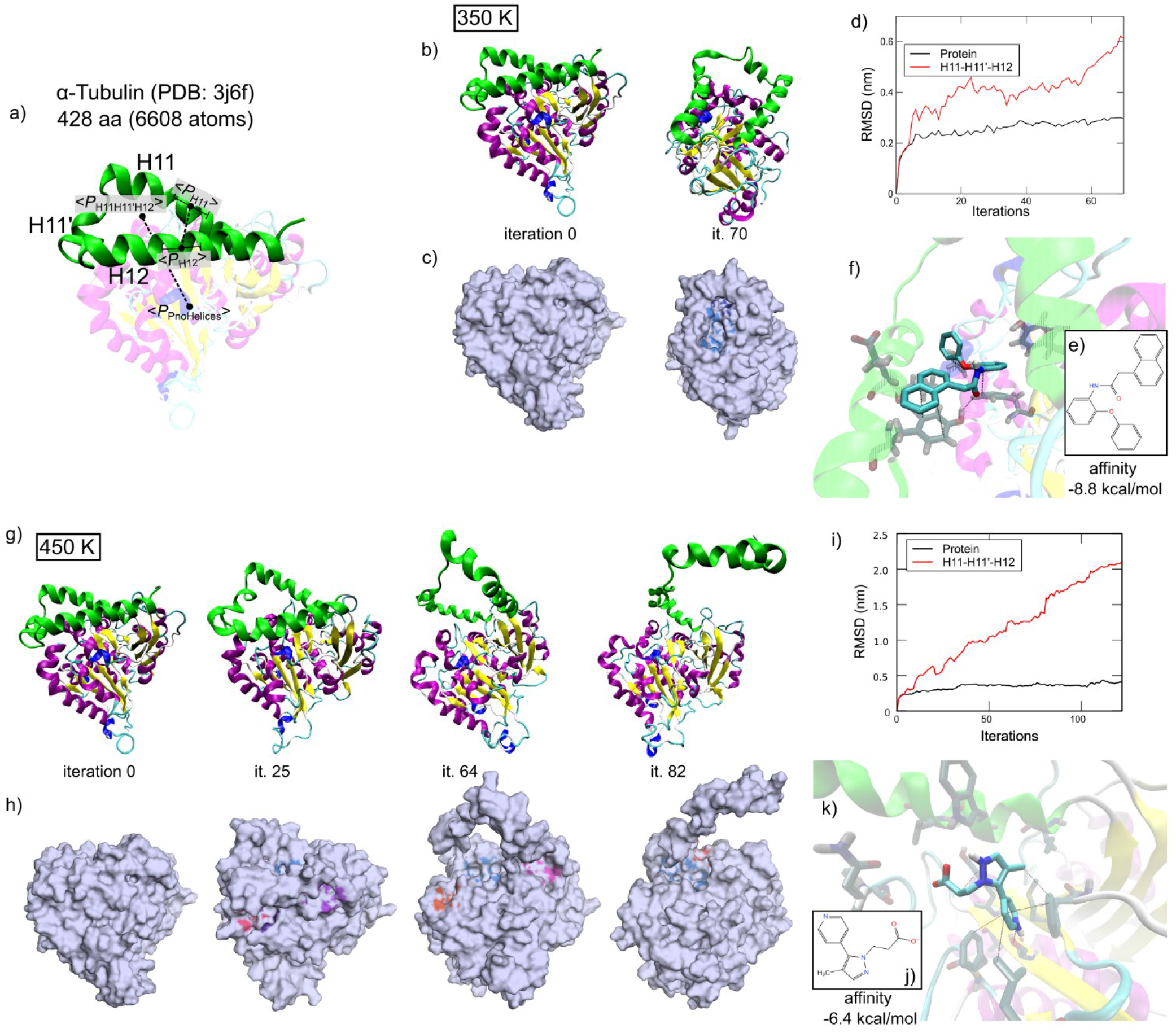
Unfolding of α-Tubulin H11-H11’-H12 domain via Discard-and-Restart MD. (a) Qualitative illustration of the CV for α-Tubulin unfolding. The large dots indicate the average coordinate point for H11, H12, the helical domain H11-H11’-H12, and the protein core (PnoHelices). b-c) Partial unfolding trajectory at 350 K and RMSD vs iterations (d). The C-terminal helical region progressively loosened up during the simulation. e) Molecule generated via Lingo3DMol on the blue pocket shown in (c). f) Pose and protein-ligand contacts, obtained using AutoDock Vina. The ligand has made 1 hydrogen bond, 2 aromatic-aromatic interactions and several hydrophobic interactions. g-h) Trajectory at 450 K and RMSD vs iterations (i). At this higher temperature, the helical domain was able to completely unfold and lose every contact with the protein core. j) Ligand generated on the light blue pocket shown in (h). k) Ligand pose and protein-ligand contacts obtained by docking simulation. The ligand has made 6 hydrophobic contacts and 1 hydrogen bond with the protein. Note that the images of the protein are not set to always show the same angle.

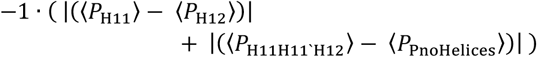

where <*P*> corresponds to the average position of the atoms of helix 11 (H11, residues 373 to 390), helix 12 (H12, residues 405 to 426), the helical domain (H11-H11’-H12, residues 373 to 426) and the rest of the protein without the helical domain (P_noHelices_, residues 1 to 372). In this way, the value of the CV decreases whenever the distance between helix 11 and helix 12 increases, or when the distance between the helical domain and the rest of the protein increases.

We want to underline that this CV for α-Tubulin unfolding does not explicitly describe the unfolded target state (which is unknown). It has been defined based on partial information available from recent experiments^65^. This means that the method is very flexible to the use of CVs, as long as they correlate with the conformational change under investigation.

### Simulation of α-Tubulin unfolding

The starting structure for the simulation was the crystal, native conformation (PDB: 3j6f). The structure has been energy minimized and equilibrated at 300K for 1 ns in implicit solvent. The simulations have been run in implicit solvent at 350 K and 450 K, one trajectory for each temperature. Each simulation required ∼1300 CPU hours.

In the trajectory computed at 350 K, the distance between helices H11 and H12 progressively increased over 70 iterations, loosening up the helical domain and exposing the residues beneath it (Figure 5b-c). The helical domain unfolded only partially and never detached from the core of the protein, because the temperature was probably too low to trigger such event (as will instead be the case for the trajectory run at 450 K). The rest of the protein underwent small changes and has been relatively stable during the simulation (RMSD < 0.3 nm, Figure 5d).

In the trajectory computed at 450 K, the helical domain unfolded completely over the course of 123 iterations, going through a complex series of transition states (Figure 5g-h). During the initial unfolding steps (iterations 0-25), the helices H11 and H12 progressively started to get further apart and the whole helical domain moved slightly up respect to the protein core, similar to the simulation at 350 K. In the successive iterations, the helical domain continued to move further up losing most of its interactions with the protein core, and H11-H11’-H12 formed a ring (iterations 25-64). Last, the C-terminal residues progressively lost all the remaining interactions, until H12 detached from the protein core and started flailing upwards. The protein core underwent small changes and has been relatively stable during the simulation (RMSD < 0.4 nm, Figure 5i). The analysis of the RMSF (Root Mean Square Fluctuation) of each atom revealed an increased structural flexibility of the H11-H11’-H12 region. This is a result of the Discard-and-Restart algorithm, which ultimately selected trajectories whose H11-H11’-H12 atoms had a slightly higher kinetic energy and were more likely to unfold. Results are shown in supplementary section 5.

### Drug discovery on the transition states

During these conformational transitions, a lot of protein regions that were previously buried by the H11-H11’-H12 domain became exposed to the solvent. We explored the possibility that these regions could have become targetable by ligands. At each iteration, we have utilized the tool P2Rank^57^, as in the previous section, in order to predict the ligand binding sites and monitor the druggability of the transitory conformations. Figure 5c and 5h show the ligand binding sites predicted for the surface regions of the transition states that were not present in the native state.

According to the study of Kuo et al.^65^, the unfolding event that we have observed is spontaneous and reversible. The drug targeting of the protein regions that are exclusively present in the transitory conformations could prevent the refolding of α-Tubulin and destabilize the microtubules, promoting the binding of Spastin to the C-terminal region. Moreover, the force required for α-Tubulin pulling out of the microtubule lattice by Spastin could be diminished in the ligand-bound state.

We have generated pocket-specific ligands using the generative AI tool Lingo3DMol^68^, a recent pocket-based 3D molecule generation method that combines language models and geometric deep learning, available as an online tool at https://sw3dmg.stonewise.cn/#/. As the ligand binding site, we have utilized the pockets which had the highest P2Rank score (i.e. the highest ligandability). The pockets that we have used are the ones highlighted in light blue in Figure 5c (second snapshot) and Figure 5h (third snapshot), respectively. Given a ligand binding site and the coordinates of the pocket center, Lingo3DMol generates a large number of drug-like synthesizable ligands (in our case 100 per pocket) that geometrically fit inside of the pocket and that have optimized non-covalent interactions. Two of the generated molecules are shown in Figure 5e and Figure 5j. However, Lingo3DMol does not provide affinity calculations and the pose of the ligand is not optimized in an energy-dependent manner. Therefore, to enhance the docking pose of the ligand in the pocket and to calculate the affinity, we have performed docking using Autodock Vina, as indicated in the previous section.

For what concerns the conformation obtained at 350 K, the docking simulation predicted a highest affinity pose of -8.8 kcal/mol, which is strong interaction. The pose of the ligand in the pocket with the main protein-ligand interactions are shown in Figure 5f.

Comparable results have been obtained with the conformation generated at 450 K. The docking simulation predicted a highest affinity pose of -6.4 kcal/mol, which is also a relatively strong interaction. Ligand pose and protein-ligand interactions are shown in Figure 5k. More details are shown in the supplementary section 6.

## CONCLUSIONS

We have presented an algorithm to speed up the MD simulation of protein conformational changes and sample the atomistic structures of the transition states, given a general CV that describes the target state. The algorithm has been benchmarked on four fast-folding proteins and on the PrP folding intermediate, and yielded results which qualitatively and quantitatively agree with previous studies. The Discard-and-Restart MD has also been employed to study the partial unfolding of the α-Tubulin C-terminal helical domain; a process that would be too time consuming by plain MD. The Discard-and-Restart MD algorithm was able to catch the unfolding process in detail and sample the conformations of the pathway.

However, everything comes at a cost. Properties such as kinetics, free energy and population states cannot be calculated from the Discard-and-Restart trajectories, because the concept of reversibility between the transition states is lost and detailed balance therefore not maintained, as the trajectories are in an out-of-equilibrium state^69^.

At each iteration of the Discard-and-Restart MD, we have monitored the druggability of the transition states by predicting the ligand binding site using AI-based tools. Conformational states with druggable local structures that would be normally hidden in the native state (i.e. cryptic pockets or folding intermediates) have been targeted for ligand docking. In this way, the Discard-and-Restart MD algorithm has the potential of speeding up the workflow of SBDD when (even partial) information about the target state is available. It strongly reduces the computational cost required for MD by up to 2000x, while maintaining the atomistic resolution and accuracy that is required by docking and protein-ligand interaction studies. We are aiming at employing this algorithm to study many more protein targets and to identify the most promising protein-ligand interactions.

## METHODS

The simulations have been performed on a Lenovo ThinkPad T470p from 2017 or on a Mac Pro from 2013. The length of the short MD simulations was chosen to be 10 ps for the fast-folding protein in implicit solvent and for PrP. The short MD runs for the fast-folding proteins in explicit solvent and for α Tubulin were instead 20ps. Starting structures have been energy minimized by steepest descent for 5000 steps or until the maximum force became lower than 5 kJ/mol/nm. Implicit solvent has been modeled using GBSA with velocity rescaling. Explicit solvent was modeled with TIP3P water, thermostatted with the Nose-Hoover thermostat and the Parrinello Rahman barostat^7,70,71^. Additional simulation details can be found in the supplementary information, section 2.

## ASSOCIATED CONTENT

### Supporting information

More information are available in the supplementary material.

## AUTHOR INFORMATION

### Author contributions

A.I. invented the Discard-and-Restart MD algorithm, designed and performed the simulations, analyzed the results and wrote the manuscript. M.G supervised the study and determined the scientific workflow.

## Acknowledgments

We thank Prof. Pietro Faccioli (University of Milan-Bicocca, Italy) and Prof. Emiliano Biasini (University of Trento, Italy) for very useful discussions and inputs without which this study would not have been possible. We also thank Jewon Im and Eddie Cavallin for programming support.

## Notes

The authors declare no competing financial interests.

