## Supplementary Information for "A Discard-and-Restart MD algorithm for the sampling of realistic protein transition states and enhance structure-based drug discovery"

**Content:**


1. **Discard-and-Restart MD and PCA model**
2. **MD simulation details and parameters**
3. **Comparison of folding CPU times of plain MD vs Discard and Restart MD**
4. **Prp (un)folding intermediate**
5. **Ligand binding analysis**
6. **References**


### Abbreviations

MD, molecular dynamics; PCA, principal component analysis; RMSD, root mean squared displacement; CV; collective variable; AI, artificial intelligence; SBDD, structure-based drug discovery; AF, AlphaFold..

### 1. Discard-and-Restart MD and PCA model

A Discard-and-Restart MD trajectory is the result of numerous short “forward” trajectories, i.e. the simulations in which at any point the CV lowers. A Discard-and-Restart folding trajectory of TrpCage in implicit solvent is shown in the figure below.

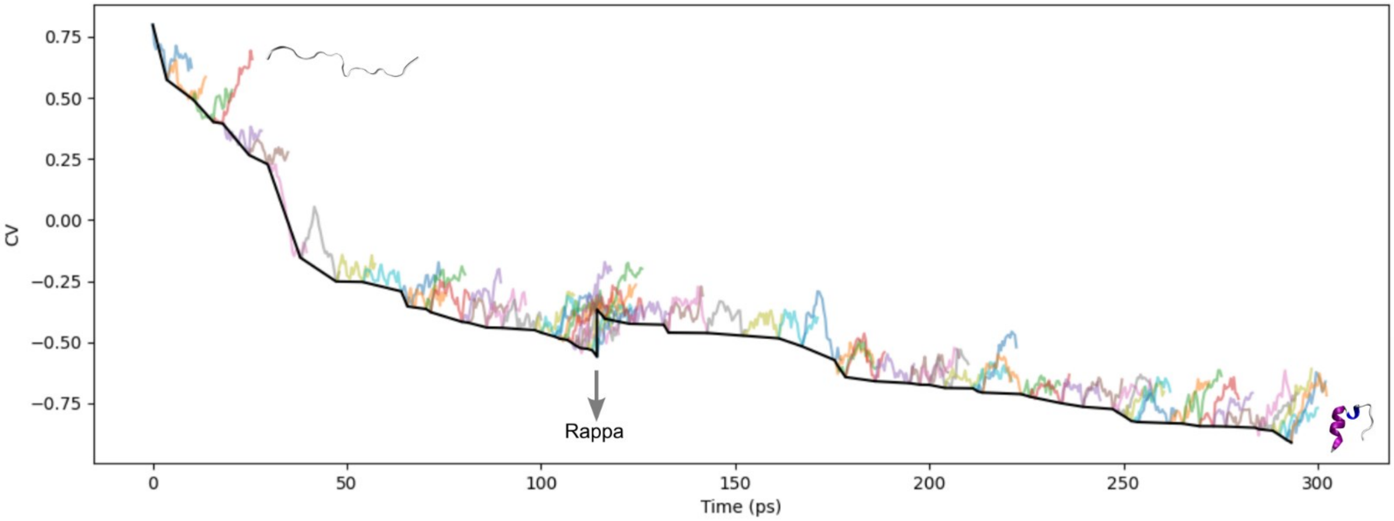


**Figure S1.1: The short forward trajectories of the Discard-and-Restart MD simulation.** The final trajectory (thick black line) is obtained by combining the conformations extracted from the short “forward” trajectories, at the timepoint where $CV_{i+1} < CV_i$ (i=iteration). Eventually, if the simulation is stuck for too many times (> *Rappa*), it is allowed to go forward regardless (grey arrow).

The PCA Eigenvectors for the calculation of the combined CV have been calculated using long MD simulation of Villin and Fip35 WW domain from DE Shaw Research[1], 100 µs of each, for a total of 396k data points. The training set used to calculate the PCA Eigenvectors therefore consisted of 396k instances and 4 features, as shown in the figure below.

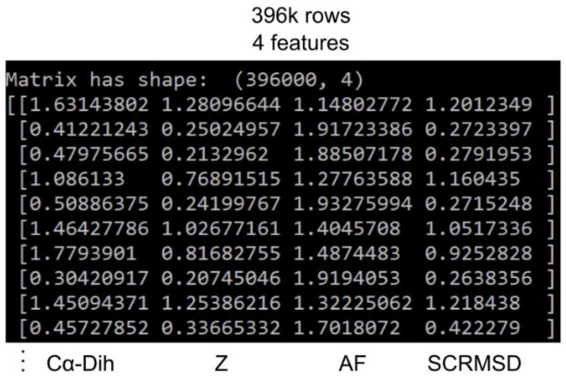


**Figure S1.2: The training set used to calculate the PCA Eigenvectors.**

### 2. MD simulation details and parameters

GROMACS mdrun .mdp file for the simulations performed in implicit solvent (at 300 K):

```
constraints          =  all-bonds
integrator           =  md
comm_mode            =  ANGULAR
nsteps               =  10000
dt                   =  0.002
```

```
pbc                  =  no
periodic_molecules   =  no
nstxout              = 200
nstvout              = 200
nstenergy            = 0
nstlog               = 0
xtc_grps             = Protein
nstxtcout            = 200
ns_type              =  grid
rlist                =  1.6
rcoulomb             =  1.6
rvdw                 =  1.6
coulombtype          =  cut-off
vdwtype              =  cut-off
cutoff-scheme        =  group
tcoupl               =  v-rescale
tau_t                =  1
ref_t                = 300.0
tc-grps              = protein
gen_vel              = yes
gen_temp             = 300.0
gen_seed             = -1
Pcoupl               = no
implicit_solvent     = gbsa
gb_algorithm         = obc
nstgbradii           = 1
rgbradii             = 1.6
gb_epsilon_solvent   = 78.3
gb_saltconc          = 0
gb_obc_alpha         = 1
gb_obc_beta          = 0.8
gb_obc_gamma         = 4.85
gb_dielectric_offset= 0.009
sa_algorithm         = ace-approximation
sa_surface_tension   = 2.092
```

Next, GROMACS mdrun .mdp file for the simulations performed in explicit solvent (300 K):

```
integrator          = md
nsteps              = 10000
dt                  = 0.002
refcoord-scaling    = all
nstxout             = 200
nstvout             = 200
nstenergy           = 0
nstlog              = 0
xtc_grps            = System
nstxtcout           = 200
continuation        = no
constraint_algorithm = lincs
constraints         = all-bonds
lincs_iter          = 1
lincs_order         = 4
ns_type             =  grid
rlist               =  1.6
rcoulomb            =  1.6
rvdw                =  1.6
nstlist             = 5
coulombtype         = PME
pme_order           = 4
fourierspacing      = 0.12
tcoupl              = Nose-Hoover
tc-grps             = Protein    Non-Protein
tau_t               = 1          1
ref_t               = 300        300
pcoupl              = Parrinello-Rahman
pcoupltype          = isotropic
tau_p               = 2.0
ref_p               = 1.0
compressibility     = 4.5e-5
```

```
pbc                  = xyz
DispCorr             = EnerPres
gen_vel              = yes
gen_temp             = 300.0
gen_seed             = -1
```

### 3. Comparison of folding CPU times of plain MD vs Discard-and-Restart MD

| Protein | Folding time[2,3] (μs) | GROMACS performance (CPU_h / ns) | Plain MD required CPU hours | D&R MD required CPU hours | Fold difference |
|---|---|---|---|---|---|
| TrpCage (implicit) | 14 | 0.5 | 6950 | 18 | 390x |
| TrpCage (TIP3P) | 14 | 18.6 | 260600 | 984 | 260x |
| Villin (implicit) | 2.8 | 1.1 | 3100 | 152 | 20x |
| Villin (explicit) | 2.8 | 7.6 | 21200 | 360 | 59x |
| Fip35 WW domain | 21 | 0.9 | 18600 | 78 | 240x |
| β-hairpin fragment | 6 | 0.5 | 3100 | 1.5 | ~2000x |

**Table S3: CPU time comparison of plain MD vs Discard and Restart MD.**

### 4. PrP (un)folding intermediate

We also performed a longer Discard-and-Restart simulation (~800 CPU hours) of PrP to continue sampling the unfolding intermediate. In 226 iterations, the domain containing the Beta-Sheet and Helix 1 separated and lost the docking contacts with the C-terminal domain of Helix 2 and Helix 3. This conformation observed in iteration 226 (Figure S4, last snapshot on the right) has also been observed in the study by Spagnolli *et al*[4] during the folding process (starting from the unfolded state).

Moreover again, the ligand binding site identified in our intermediate conformations (Figure S4) matches with the one identified in the original study by Spagnolli *et al*. (which involved the residues 152, 153, 156, 157, 158, 187, 196, 197, 198, 202, 203, 205, 206, 209). In our case, the pocket identified in the conformation of iteration 89 involves the residues 159, 160, 187, 190, 191, 198, 202, 206, and the location of the pocket is substantially identical (data not shown).

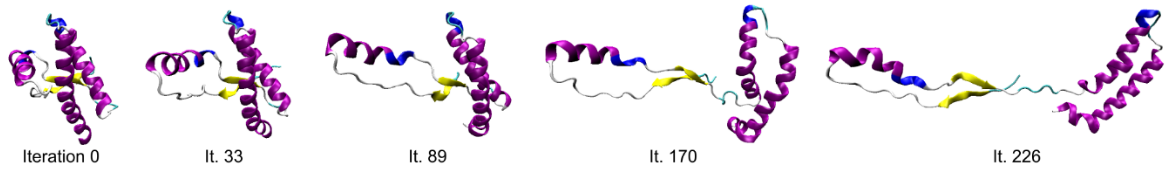


**Figure S4: Longer Discard-and-Restart simulation of the PrP unfolding intermediate.**

### 5. Flexibility of the Discard-and-Restart structures

To characterize the flexibility of the structures during the Discard-and-Restart simulations, we have calculated the RMSF (Root Mean Square Fluctuation) for each atom of the protein. The figure below shows the RMSF for α-Tubulin and is compared to the RMSF of standard MD simulations. It is possible to see that the RMSF is pretty much identical between standard MD (red) and Discard-and-Restart MD (blue), except for the part regarding the helical domain H11-H11’-H12, which was part of the CV loss function. This difference probably arises from the selection process of the Discard-and-Restart algorithm, which has been selecting the trajectories that had a higher initial velocity of the atoms in the H11-H11’-H12 region (atoms from 5701 to 6608), therefore more likely to unfold and decrease the CV loss function.

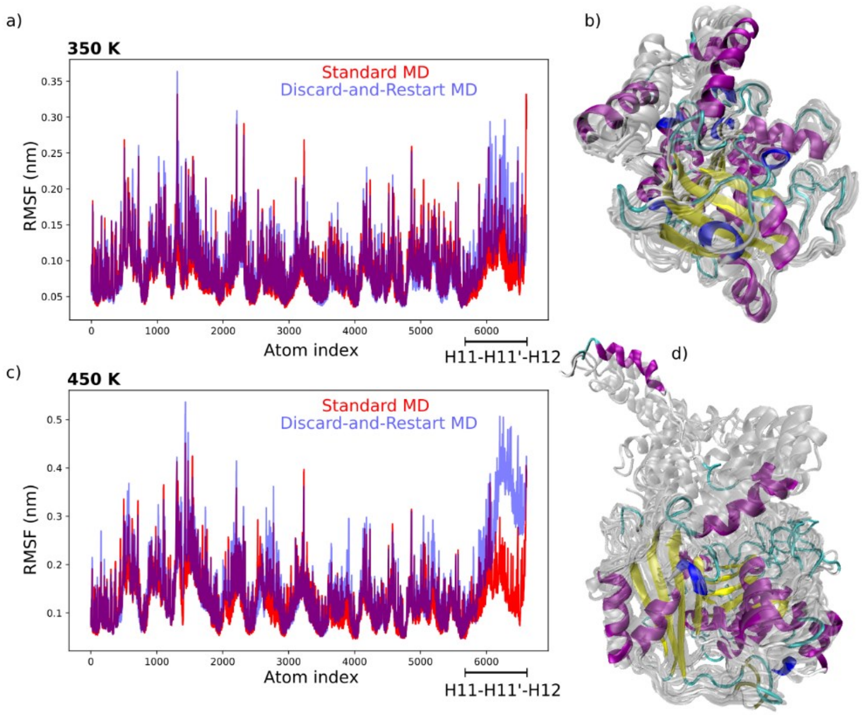


**Figure S5: Characterization of the RMSF of Discard-and-Restart compared to standard MD simulations.** a) RMSF for each atom of α-Tubulin for the system at 350K, and overlay of the structures of the H11-H11’-H12 unfolding simulation (b). c-d) Same but for the system studied at 450K.

### 6. Ligand binding analysis

Lingo3DMol generates a multitude of molecules that geometrically fit into the pocket[5]. The molecules are not scored by affinity or any other docking-related metrics, so the choice at this point remains arbitrary. Additional examples of ligands generated for the pocket of Figure 4h (blue pocket) are shown below.

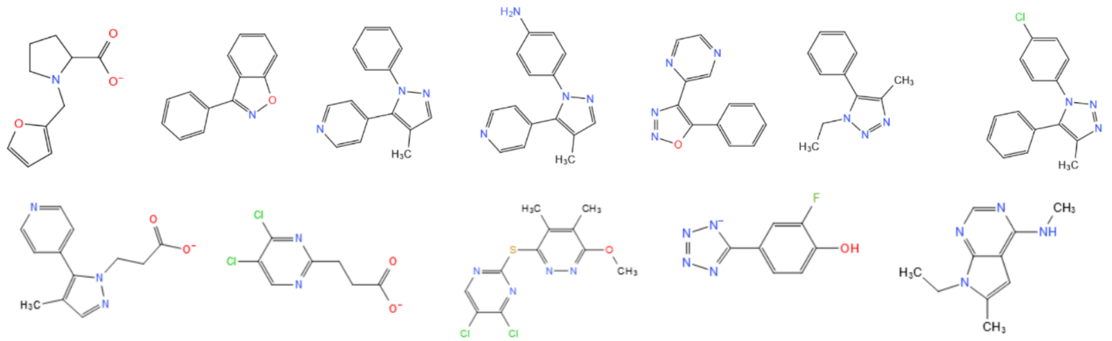

**Figure S6.1: Twelve pocket-specific molecules generated by Lingo3DMol.**

The molecule of choice must then be docked into the pocket via docking simulation, for example using AutoDock Vina as we have used in our study[6,7]. There are then additional tools that help in the visualization of the protein-ligand interactions, such as ProteinsPlus[8] . It shows the main contacts (e.g. hydrophobic, aromatic, hydrogen bonds...) between the protein and the ligand.

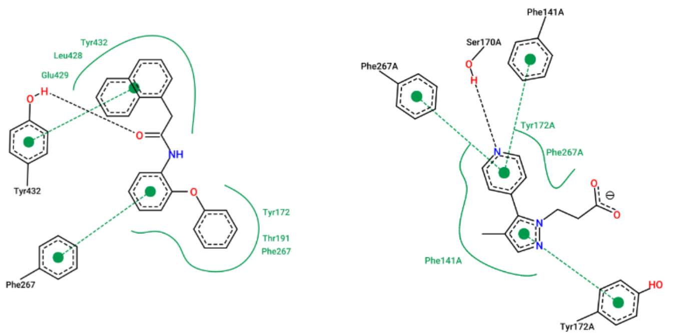


**Figure S5.2: Ligand-protein binding analysis using ProteinsPlus.** Interaction analysis for the ligand docked in the pocket shown in Figure 45 (left) and 5k (right).
